# Spatial epidemiology of networked metapopulation: An overview

**DOI:** 10.1101/003889

**Authors:** Lin Wang, Xiang Li

**Affiliations:** Adaptive Networks and Control Laboratory, Department of Electronic Engineering, Fudan University, Shanghai 200433, China; Centre for Chaos and Complex Networks, Department of Electronic Engineering, City University of Hong Kong, Hong Kong SAR, China

**Keywords:** Complex networks, Epidemiology, Spatial dynamics, Metapopulation

## Abstract

An emerging disease is one infectious epidemic caused by a newly transmissible pathogen, which has either appeared for the first time or already existed in human populations, having the capacity to increase rapidly in incidence as well as geographic range. Adapting to human immune system, emerging diseases may trigger large-scale pandemic spreading, such as the transnational spreading of SARS, the global outbreak of A(H1N1), and the recent potential invasion of avian influenza A(H7N9). To study the dynamics mediating the transmission of emerging diseases, spatial epidemiology of networked metapopulation provides a valuable modeling framework, which takes spatially distributed factors into consideration. This review elaborates the latest progresses on the spatial metapopulation dynamics, discusses empirical and theoretical findings that verify the validity of networked metapopulations, and the application in evaluating the effectiveness of disease intervention strategies as well.

## 1 Introduction

The term metapopulation was coined by Levins [1] in 1969 to describe a population dynamics model of insect pests in farmlands, yet the perspective has been broadly applied to study the effect of spatially distributed factors on evolutionary dynamics [2], including genetic drift, pattern formation, extinction and recolonization, etc. The development of metapopulation theory, in conjunction with the fast development of complex networks theory, lead to the innovative application of the networked metapopulation in modeling large-scale spatial transmission of emerging diseases. This interdisciplinary research field has attracted much attention by the scientific communities from diverse disciplines, such as public health, mathematical biology, statistical physics, information science, sociology, and complexity science. New insights are contributed to understanding the spatial dynamics of epidemic spreading, which provides valuable support to public healthcare.

This review presents a survey of recent advances in the emergent discipline of networked metapopulation epidemiology, which is organized as follows. Section 2 introduces some preliminaries of the compartment model, network epidemiology, and networked metapopulation, and also elucidates their relevance. Section 3 specifies the validity of networked metapopulation. Section 4 focuses on the recent progresses on metapopulation dynamics. The application in evaluating the performance of intervention strategies is presented in Section 5, and some outlooks are provided at last.

## 2 Dynamical models of infectious diseases: From single population to networked metapopulation

### 2.1 Compartment model

To study the phenomena of epidemic spreading in human society, a variety of dynamical models have been proposed [3,4]. The compartment model is one of the simplest yet basic epidemic models, which was first introduced by Bernoulli [5] in the 18th century. Assuming that a population of individuals is mixed homogeneously, this model organizes the persons into different compartments (states), according to their health status, e.g., susceptible (denoted by S, those healthy ones who may acquire the infection), infectious (I, those infected ones who are contagious), and recovered (R, those who are recovered from the disease). Within each compartment, all individuals are identical. The transitions between different compartments depend on the specific transition rates. For example, the transmission rate *β* represents the infection probability for a susceptible individual that encounters an infectious person, and the recovery rate *μ* represents the probability with which an infectious individual is recovered.

If the disease could not endow recovered persons with a long lasting immunity but infect them again, e.g., seasonal flu, asthma, gonorrhoea, the related epidemic reactions are well described by the so called SIS model; otherwise, if recovered people become immune permanently to the disease, e.g., pandemic influenza, pertussis, smallpox, the epidemic dynamics can be characterized by the SIR model properly. Figures 1(a)-(b) illustrate the relevant compartment transitions in the SIS and SIR models, respectively. The dynamical evolution of these models can be simply delineated by ordinary differential equations [3].

**Fig. 1.**
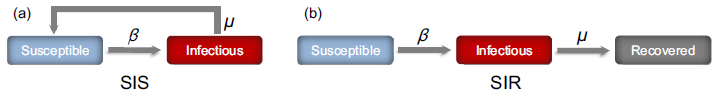
(*Color online*) Schematic illustrations of the SIS (a) and the SIR (b) compartment models, where *β*, *μ* denote the transmission rate and the recovery rate, respectively.

One key parameter characterizing the severity of a disease is the basic reproductive number, *R*_0_, which identifies the expected number of infected individuals generated by introducing an infectious carrier into an entire susceptible population. This parameter signifies the epidemic threshold applied for predicting whether or not an infectious disease will prevail. Typically, given a “well-mixed” population, *R*_0_ = *β*/*μ*. If *R*_0_ < 1, the disease dies out quickly, which implies that the population remains at the disease-free state.

### 2.2 Network epidemiology

Due to the ubiquity of complex systems in modern society, the study of complex networks becomes prosperous [6–9]. The Internet and human friendship networks are just a few examples that can be regarded as systems comprised of a large number of connected dynamical units. The most intuitive approach of modeling such complex systems is to treat them as networks, where nodes represent component units and edges represent connectivity. Importantly, empirical findings have unraveled the presence of universal features in most socio-technical networks, e.g., small-world [10], scale-free (SF) [11], which inspires extensive studies towards a better understanding about the impact of population infrastructures (network connectivity) on dynamical processes [12–15], including robustness [16,17], synchronization [18–20], consensus [21–24], control [25–28], evolutionary game [29–36], traffic routing [37–39], selforganized criticality [40–43], etc.

Assuming that interactive individuals are mixed homogeneously, the aforementioned epidemic compartment model neglects the significance of population connectivity. Such simplification can hardly solve new puzzles emerged in the present networking society. For example, why is it extremely difficult to eradicate computer viruses from the Internet or the World Wide Web, and why do those viruses have an unusual long lifetime [44]? Similar matters have been observed in diverse systems, ranging from the web of human sexual relations to vaccination campaigns [4]. One key factor inducing such problems is the scale-free property of the networked systems, which causes a serious trouble that the threshold of disease outbreak vanishes [45]. Within complex networks, the basic reproductive number is 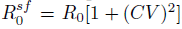, with *CV* identifying the coefficient of variation of the degree distribution (degree represents the number of edges *k* per node) [46]. For large networks taking on a scalefree heterogeneous topology, *R*_0_ is always larger than 1 no matter how small the transmission rate may be, due to the infinite variance of the degree distribution.

This meaningful finding has motivated the research of network epidemiology, which concerns particularly the spreading of epidemics in human social networks [4,14,47]. Many subsequent works investigated extensively the epidemic threshold on networks with special topological features, such as degree correlations [48], small world [49], community [50], edge length [51], and K-core [52]. [53,54] demonstrated that the vanishing epidemic threshold of the SIS model derives from the active behavior of the largest hub, which acts as a self-sustained source of the infection. Such disastrous effect of highly connected hubs can also be observed in reality, such as the presence of core groups in the propagation of sexually transmitted diseases, and the appearance of the patient zero that induces the dissemination of human immunodeficiency virus (HIV). Considering that the threshold condition generally predicts the final state of the epidemic evolution, Li and Wang [49] studied the relaxation behavior of epidemic spreading before reaching a final disease-free or endemic phase.

### 2.3 Networked metapopulation

Although the performance of public healthcare systems has been improved prominently to weaken the threat of emerging diseases, it is impossible to entail a world free of infectious pathogens [55]. From the beginning of this new century, we have already witnessed several cases of the large-scale geographic transmission of pandemics. In 2003, through the international airline network, the SARS coronavirus (SARS-CoV) was rapidly transmitted from Hong Kong to more than 30 countries [56,57]. Several years later, in 2009, the A(H1N1) swept across the world through public transportation networks again: With only 3 to 4 months, it had spread over about 200 countries [58–61]. Recent potential invasion of avian influenza A(H7N9) poses a new challenge [62–65]. It seems that the widespread risk of emerging diseases is higher than before.

This urgent circumstance stems from the changes of human social ecology in population distribution as well as human mobility patterns [66,67]. Crowded metropolises resulting from the urbanization process induce people’s frequent contacts, and the fast development of massive transportation (e.g., civil aviation) generates a nonlocal pattern of human mobility, sharply reducing the time of travel as well as the distance between populous cities.

It is not convincing to describe the large-scale spatial pandemic spreading by directly following the routine of network epidemiology, since the network perspective still concerns the epidemic outbreak in a single population, despite considering the connectivity structure among hosts. This can hardly capture the key features of spatial transmission of infectious diseases: epidemics prevails inside separate locations such as cities, each of which can be regarded as a population, and is transmitted among populations through the travel of infected individuals.

Spatial distribution of populations and human mobility among connected locations are the pivotal elements mediating the transmission of pandemic diseases. To introduce spatially distributed factors into modeling substrates, it is intuitive to generalize the network model by defining each node as a subpopulation that has a specific location, in which a population of individuals interplays according to the compartment rule. People are also permitted to transfer among subpopulations through mobility networks. This individual-network frame organizes the entire system into networked populations, leading to an important class of model in modern epidemiology, namely, the networked metapopulation. Figure 2 illustrates the basic modeling structure.

**Fig. 2.**
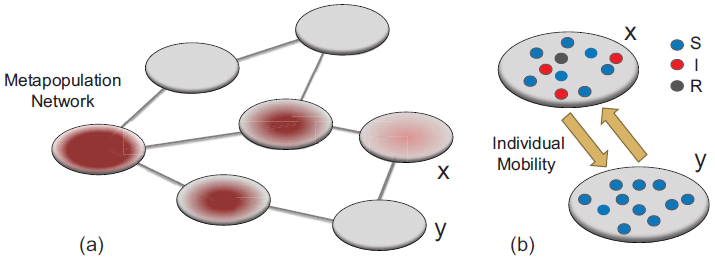
(*Color online*) Illustration of the individual-network frame of the networked metapopulation model. **a** The model is composed of a network of subpopulations. The disease transmission among subpopulations stems from the mobility of infected individuals. **b** Each subpopulation refers to a location, in which a population of individuals interplays according to the compartment rule (e.g., SIR) that induces local disease outbreaks. Individuals are transferred among subpopulations via mobility networks.

## 3 Validity of networked metapopulation

Aside from the above conceptual descriptions, it is also essential to verify the validity of the model from theoretical as well as empirical perspectives.

Developing a probabilistic metapopulation with the consideration of long-range human migrations via a worldwide aviation network composed of 500 largest airports, Hufnagel et al. [68] first demonstrated the feasibility of forecasting the real-world transmission of SARS through computational approaches. To study the spatiotemporal patterns of the transmission process, Colizza et al. [69] defined a statistical measure based on the information entropy, which quantifies the disorder level encoded in the evolution profiles of disease prevalence. Comparing the pandemic spreading on a data-driven networked metapopulation with that on random reshuffled models providing null hypotheses, the authors unveiled the presence of a high-level heterogeneity in the geographic transmission of epidemics.

To assess the predictability of metapopulation models, one typical approach focuses on the coincident extent between the simulation results and the realistic surveillance reports for each contaminated region, which is an arduous task due to the sophisticated calibration of parameters as well as the unavoidable noise presented in the surveillance process. Concerning the logistical feasibility of the model, one can resort to an alternative simple means of inspecting the evolution of related scaling laws [70], which is relevant to critical transition patterns. The scaling theory concerns the functional relations describing the data collapsing onto a power-law curve, and the relations of the critical-point exponents [71].

The Zipf’s law and the Heaps’ law are two representative scaling laws that usually emerge together in various complex systems, however, their joint emergence has hardly been clarified [72]. Using the data of laboratory confirmed cases of SARS, H5N1, and A(H1N1) to analyze the joint emergence of these two scalings in the evolution process of large-scale geographic transmissions, Wang et al. [70] unraveled a universal feature that the Zipf’s law and the Heaps’ law are naturally shaped to coexist at the initial stage of an outbreak, while a crossover comes with their incoherence later before reaching a stable state, where the Heaps’ law still presents with the wane of the strict Zipf’s law. With the census populations and domestic air transportation data of the United States (US) [73,74], a data-driven metapopulation network model on the US country level is developed to analyze the evolution patterns of scaling emergence. In contrast with a random reshuffled model with a homogeneous structure, the data-driven heterogeneous metapopulation successfully reproduced the scaling transitions observed in the real-world pandemics. This demonstrates that the high-level heterogeneity of infrastructure plays a key role in characterizing the spatial transmission of infectious diseases, which also provides a new insight to clarifying the interdependence between the Zipf’s and Heaps’ scaling laws.

Within each subpopulation, the individuals are mixed homogeneously, according to the coarse-grained approximation of the metapopulation framework. Interestingly, this assumption can be supported by recent empirical studies on the intra-urban human mobility. The analysis of the data generated by the mobile phone or GPS shows that human movement in the urban scale (e.g., inside a city) generally has an exponential or binomial trip-length distribution [75–79]. Although this does not simply mean that short-range human mobility is random, the related dynamical feature is similar with that of the Boltzmann gas, if the relevance among individuals is so weak as to be negligible [76,80,81]. Accordingly, the homogeneous mixing (within each subpopulation) assumption is adopted to ease the computation.

More promisingly, full-scales computational models become increasingly popular, due to the continuous increase of computer power as well as the fast technical developments of data collection and processing [82–84]. In some cases, the real-time forecast of pandemic spreading is becoming reality [85]. Technical details for the estimation and validation of a large number of parameters in these models are beyond the interest of this review. Next section focuses on the recent theoretical progress of metapopulation dynamics.

## 4 Two scales of dynamics: Recent progress

As stated in Section 2, the networked metapopulation model is constructed with the individual-network frame, where the individuals are organized into social units (e.g., villages, towns, cities) defined as subpopulations, which are connected by transportation networks that identify the mobility routes. The disease prevails inside each subpopulation due to interpersonal contacts, and is transmitted among subpopulations through the mobility of infected individuals. Typically, the model is comprised of two scales of dynamics: (i) disease invasion among different subpopulations; (ii) disease reaction within each subpopulation. Recent progresses on these two aspects are specified here.

### 2.1 Inter-subpopulation invasion

The substrate of metapopulation depends on the spatial structure of social environment, such as transport infrastructures and mobility patterns. The lack of fine-grained data capturing structural features of human mobility systems leads to the traditional application of random graphs or regular lattices, which assumes homogeneous infrastructures for the mobility substrates. To generalize metapopulation models with network approaches, the first attempt was contributed by Rvachev and Longini [86], in which 52 major cities worldwide in that epoch were connected through an intercity aviation transportation network. They applied this mathematical model to simulate the global spread of the 1968–1969 Hong Kong (H3N2) flu.

Subsequently, comparing the effect of non-local human anomalous diffusion with that of the ordinary diffusion behavior, Brockmann et al. unraveled that long-range human mobility and interactions generate novel irregular spreading patterns without an apparent wavefront [87]. Such complex dynamical features require a mathematical description of fractional diffusion equations, and they are also well captured by the networked metapopulations. Colizza et al. [69] developed a global stochastic metapopulation model in, using the data of worldwide scheduled flights and census populations to establish a complete worldwide air transportation network (more than 3000 airports). They studied the predictability and the reliability of the pandemic forecast with respect to the intrinsic stochasticity, and declared that the topological heterogeneity reduces the predictability, whereas the high-level heterogeneity of traffic flows improves the pandemic predictability.

As illustrated by Fig. 3 **a**, air traffic network acts as a major channel serving human long-range travels, which mediates the pandemic transmission on a large geographic scale. The epidemic dynamics occurred under this scenario is well characterized by the reaction-diffusion processes [88], which are also widely applied to model phenomena as diverse as genetic drift, chemical reactions, and population evolution [2].

**Fig. 3.**
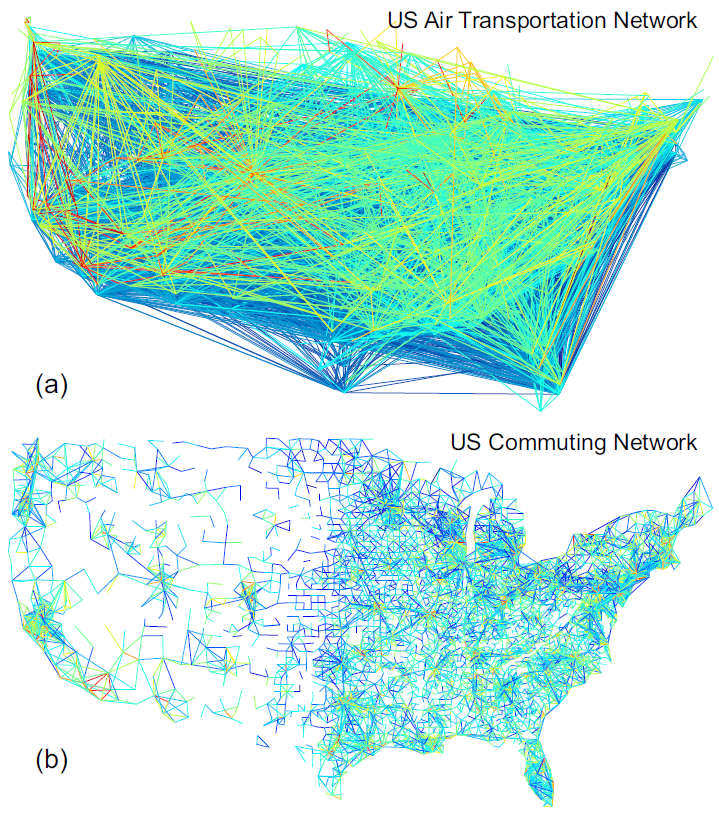
(*Color online*) Air transportation network (a) vs. commuting network (b) of the US. Long-range airlines dominate the air transportation network, whereas the commuting routes are much geographically localized.

From a theoretical viewpoint, it is significant to analyze the epidemic threshold, which is instructive for the assessment of the disease transmissibility as well as the outbreak potential. Such information is also important to regulate the implementation of intervention strategies. Based on the empirical evidences that the topology of various socio-technical networks including the airline network presents a high-level heterogeneity, Colizza et al. [88] studied the effect of general heterogeneous networks, demonstrating that the epidemic threshold is significantly decreased with the augmentation of topological fluctuations. Considering that the theory developed in Colizza et al. [88] is based on the simplification that individual diffusion rate per subpopulation is inversely proportional to the degree of subpopulations, Colizza and Vespignani [89] generalized the study by introducing more realistic diffusion rules, such as the traffic- and the population-dependent patterns. Importantly, using the approach of branching process, Colizza and Vespignani [89] proposed a global invasion threshold, *R*_*_, which distinguishes the lower bound condition for transmitting the infections to downstream unaffected subpopulations. The formula of *R*_*_ can be summarized as 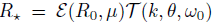, which combines the epidemiology factors *ε* (*R*_0_, *μ*) with the diffusion properties of mobility networks 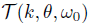. For large networks with a high-level topological heterogeneity, the mobility item 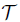 diverges, 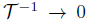, thus *R*_*_ is always larger than unity, which leads to a decreased epidemic threshold. Based on the observation that human beings usually do not perform random walks, yet have specific travel destinations, Tang et al. [90] addressed the effect of objective traveling behavior which enlarges the final morbidity.

The above studies mainly concern the influence of human random diffusion, usually defining the mobility scheme as a Markovian memoryless diffusive process [91]. Recent empirical findings on human mobility have shown the crucial role of commuting mobility in human daily transportation, which is reflected by the individual recurrent movement between frequently visited locations such as household, school, and workplace [92–95]. Fig. 3**b** visualizes the US commuting network with the census data on commuting trips between counties [96]. Evidently, the structural features are different between the commuting network and the air transportation network.

It might be infeasible to analyze the non-Markovian properties of human commuting with previous reaction-diffusion theory. In this regard, Balcan and Vespignani [91] extended the metapopulation framework by considering the impact of human recurrent commuting, which assumes that individuals remember their subpopulations of residence, with a constraint that commuters staying at their destination subpopulations cannot continue moving to other places but return to the residences with a certain rate. The approach of timescale separation is applied to perform theoretical analysis, since in reality the number of frequent commuters only accounts for a small fraction of local populations. This leads to a mean-field description of stationary populations distribution. Generalizing the theory of branching process, Balcan and Vespignani [97] obtained the global invasion threshold for the reaction-commuting networked metapopulation systems, which establishes a new threshold relevant to the typical visiting duration of commuters. With a high return rate, the sojourn time (i.e., length of stay) of infected commuters might be too short to transmit the infection to susceptibles in adjacent unaffected subpopulations.

To study the dynamical differences between the reaction-commuting and the reaction-diffusion processes, Belik et al. [98] analyzed their respective traveling wave solutions on the one dimensional lattice. As the diffusion rate increases, spatially constrained human commuting generates a saturated threshold of the wave front velocity, whereas the reaction-diffusion model has an unbounded front velocity threshold. Such distinction implies that the estimation of transmission speed might be overestimated under the reaction-diffusion framework. Besides, they have also found that the characteristic sojourn time spent by commuters induces a novel epidemic threshold. Since airline traffic and ground commuting networks both serve human routine transportation, Balcan et al. [94] developed a multiscale networked metapopulation model, where the commuting networks in about 30 countries were embedded into the worldwide long-range air transportation network. The introduction of short-range commuting mobility enhances the synchronization of epidemic evolution profiles for subpopulations in close geographical proximity.

Human beings are intelligent. Their risk perception and adaptive abilities promote the active response to epidemic outbreaks, which might in turn alter the disease propagation [99–101]. Many works [102–111] have investigated the effect of disease-behavior mutual feedback on compartment models as well as network epidemiology, and recent research topics also begin the generalization to deal with human behavior of mobility response. For example, [112,113] analyzed the impact of self-initiated mobility on the invasion threshold, showing a counterintuitive phenomenon that the mobility change of avoiding infected locations with high prevalences enhances the disease spreading to the entire system.

### 2.2 Intra-subpopulation contagion

The above studies focus on understanding the influence of inter-subpopulation human mobility patterns, generally assuming that the individuals behave identically in each subpopulation. However, the diversity of individual behaviors in different subpopulations also affects the pandemic spreading.

Although it is well-known that human contacts have crucial impact on the spatiotemporal dynamics of infectious diseases in a population [3], previous works assumed that individual contact patterns are identical among all subpopulations. Since the basic reproductive number, *R*_0_, is equivalent to the same constant in all subpopulations, it is predictable that the epidemic attack rates as well as evolution profiles in different areas are similar, as one can clearly observe in [114].

At the intra-subpopulation scale, aside from the empirical support from the data analysis of intra-urban human mobility (see Section 3), the feasibility of the “well-mixed” contacts assumption is also consistent with the recent findings on interactive patterns of human contact. For example, diverse digital instruments, e.g., wireless sensors [115], active Radio Frequency Identification (RFID) devices [116,117], and WiFi [118–120] (we resort to the WiFi technology in our social experiments, due to its ubiquity in urban areas), have been deployed in realistic social circumstances to collect the data of human close proximity contacts [121]. The data analyses have unveiled an unexpected feature that the squared coefficient of variance is quite small for the distribution of the number of distinct persons each individual encounters per day [115–119], which implies the presence of a characteristic contact rate within each subpopulation.

Note that the characteristic contact rate might vary evidently in different subpopulations. As illustrated by empirical studies [122,123], in reality, location-specific factors are the potential drivers resulting in a substantial variation of disease incidences between populations. Inspired by this finding, Wang et al. [124,125] introduced two categories of location-specific human contact patterns into a phenomenological reaction-commuting metapopulation model. A simple destination-driven scenario is considered first, where individual contact features are determined by the visited locations. Since the residence and the destination can be distinguished by the commuting mobility, an origin-driven scenario is also introduced, where the contacts of individuals are relevant to their subpopulations of residence. Figures 4(a)-(b) illustrate the modeling structures of these two scenarios.

**Fig. 4.**
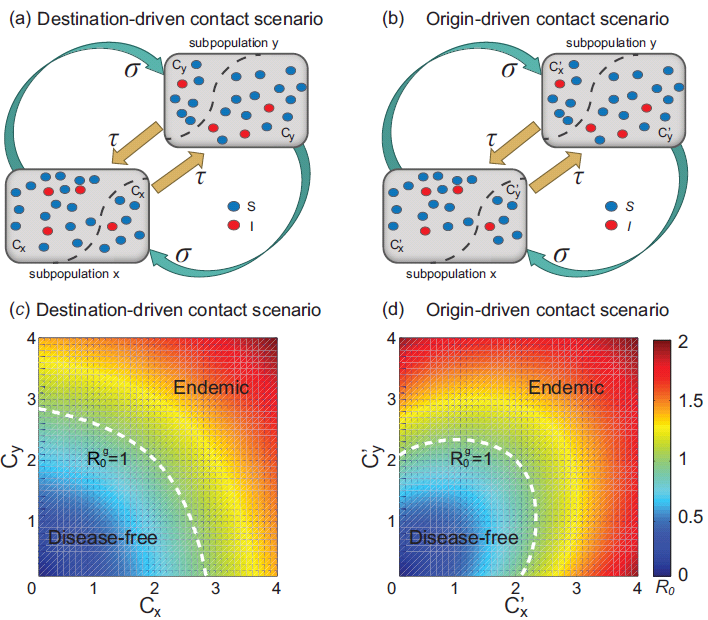
(*Color online*) Effect of location-specific human contact patterns. **a**–**b** illustrate the structure of the phenomenological metapopulation model used in [124], where the reaction-commuting processes couple two typical subpopulations *x*,*y*. In the destination-driven scenario (a), individual characteristic contact rates (*c*_*x*_,*c*_*y*_) depend on the visited locations, while in the origin-driven scenario (b), the contacts of individuals correlate to their subpopulations of residence. **c**–**d** present the phase diagrams of the global 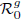 under these two scenarios, respectively. The white dashed curve in each panel shows the global threshold 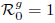 obtained through the NGM analysis. From Ref. [124].

In these cases, it is infeasible to analyze the invasion threshold through the theory of branching process, since the prerequisite of identical basic reproductive number in all subpopulations is invalid. Instead, the next generation matrix (NGM) approach [126] can be applied to analyze the global outbreak threshold 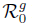 here. Due to the mixing of individuals with heterogeneous contact capacities in each subpopulation, which is analogous to the effect induced by annealed heterogeneous networks [45], the addressed location-specific contact patterns reduce the epidemic threshold significantly, and thus favor disease outbreaks in contrast to the traditional homogeneous cases. Figs. 4 **c**–**d** show the phase diagrams of the global 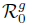 under these two types of contact patterns, respectively. Interestingly, the variance of disease prevalence under the destination-driven scenario has a monotonic dependence on the characteristic contact rates, whereas under the origin-driven scenario, counterintuitively, the increase of contact rates weakens the disease prevalence in some parametric ranges. This topic was also extended to study the metapopulation network, which unraveled a new problem of disease localization, i.e., the epidemic might be localized on a finite number of highly connected hubs.

Other types of human behavioral diversity have also been considered recently. Motivated by the evidence that the diversity of travel habits or trip durations might yield heterogeneity in the sojourn time spent at destinations, Poletto et al. [127] studied the impact of large fluctuations of visiting durations on the epidemic threshold, finding that the positively-correlated and the negatively-correlated degree-based staying durations lead to distinct invasion paths to global outbreaks. Based on the observation that the specific curing (recovery) condition depends on the available medical resources supplied by local health sectors, Shen et al. [128] studied the effect of degree-dependent curing rates, which demonstrates that an optimal intervention performance with the largest epidemic threshold is obtained by designing the heterogeneous distribution of curing rates as a superlinear mode. Since the epidemic spreading is also relevant to casual contacts during public gatherings, Cao et al. [129] introduced the rendezvous effect into a bipartite metapopulation network, and showed that the rendezvous-induced transmission accelerates the pandemic outbreaks.

## 5 Performance of intervention strategies

The study of metapopulation model not only expands our knowledge on the dynamics of spatial epidemic spreading, but also manifests the power in evaluating the performance of intervention strategies. For example, although the strategy of travel ban is usually deployed during a pandemic outbreak in reality, it is unclear whether the effectiveness is excellent enough in limiting the pandemic spreading. Counterintuitively, recent studies have unraveled the limited utility of travel restrictions: Even if the worldwide air traffic is decreased to an unprecedented low level, e.g., less than 10 %, the disease landing to unaffected regions is only postponed several weeks [130–133]; the contribution to reducing the morbity is also quite limited [130,131,134]. Such findings are consistent with the aforementioned fact that the global invasion threshold is decreased significantly by the presence of the high-level topological heterogeneity.

It thus becomes urgent to study the controllability of intra-subpopulation measures, such as the usage of vaccine or antiviral drugs, and the implementation of community-based interventions, which are typical containment strategies suggested by the World Health Organization (WHO) [55]. To estimate and also to improve the performance of disease response plans on decreasing the morbidity, large-scale computational simulations have been performed extensively to study various types of pharmaceutical interventions [4,14,56,57,60,68,134–139], which aid in identifying the targeted-groups and guiding the deployment of limited resources.

Despite technical difficulties, it is probable to analyze the delaying effect of different strategies. With the theory of renewal process, Wang et al. [140] developed a general mathematical framework to deal with the scenario of minimum metapopulation, where two typical subpopulations are connected by the travel flows. This is a rational approximation of the initial stage of an outbreak. It is shown that with a short response time, the intra-subpopulation measures perform much better than that of the inter-subpopulation travel restrictions. However, this advantage is weakened considerably as the response time increases.

Recent clinical evidences obtained from the real-world pandemic campaigns have uncovered new problems on the prompt response with pharmaceutical interventions. For example, there presents an unavoidable delay of 4–6 months for developing the proper vaccine against a particular pandemic virus [141–143]; and an extensive usage of antiviral drugs might induce the prevalence of antiviral resistance [144–146]. Therefore, it is crucial to thoroughly examine the effectiveness of community-based interventions by using the models of networked metapopulation, which deserves more efforts in near future.

## 6 Conclusions & outlooks

Networked metapopulation contributes an ideal epidemic modeling platform, which promotes our understanding on the dynamics of large-scale geographic transmission of emergent diseases. The models have the potential to be applied in the real-time numerical pandemic forecast, and are also very useful in evaluating the effectiveness of disease response strategies.

Recently, the good, the bad and the ugly facts of the Big Data have triggered extensive debates around the world. The interdisciplinary research of metapopulation epidemiology establishes a paradigm for the study of data science, since one remarkable progress in this field is the innovative usage of fine-grained data in verifying key assumptions and in establishing model substrates. Technical developments in the data collection, processing and analysis not only offer key insights into the dynamical properties of human mobility infrastructures as well as human behavioral diversity, but also raise new questions referring to their influences on the spatial transmission of emerging infectious diseases. Such methodology can be applied to study diverse types of contagion phenomena, including the spreading of computer viruses, information, innovations, emotion, behavior, crisis, culture, etc.

At the end of discussions, some open questions still deserve to be addressed. The development of the sophisticated computational techniques and the consideration of detailed human/population dynamics are quite important for the research of spatial epidemiology. However, it is also crucial to understand the fundamental principals governing the complex contagion phenomena [147]. In this regard, an interesting question poses itself, namely, whether it is possible to define a unified mathematical framework that can characterize different kinds of spatial dynamics models of emerging diseases.

It is also probable to generalize present theoretical results to deal with reverse problems, such as the identification of infection sources [147–149], possible mobility networks [150], and disease invasion process. Such inference problems are valuable to establish an optimal response plan for tracing and preventing the pandemics.

## Acknowledgements

We appreciate the two anonymous referees for their valuable comments. We are grateful to the instructive discussions with Guanrong Chen, Joseph T. Wu, Shlomo Havlin, Ming Tang, Daqing Li, Xiaoyong Yan, Zhen Wang, Jianbo Wang, Lang Cao, Xun Li. We thank Yan Zhang for the help of preparing the figures. We acknowledge the support from the National Basic Research Program (2010CB731403), the National Natural Science Foundation (61273223), the Research Fund for the Doctoral Program of Higher Education (20120071110029) of China, and the Hong Kong Research Grants Council under the GRF Grant CityU 1109/12. LW also acknowledges the partial support by Fudan University Excellent Doctoral Research Program (985 Project).

